# Bone and cartilage staining protocol for large avian specimens: late-stage embryonic emu (*Dromaius novaehollandiae*) and adult chickens (*Gallus gallus*)

**DOI:** 10.1101/2025.03.24.645048

**Authors:** Meredith J. Taylor, Todd L. Green, Gabby Guilhon, Talia Lowi-Merri, Akinobu Watanabe

## Abstract

The ability to consistently and clearly visualize discrete anatomical parts of a specimen is crucial for biological research. For skeletal tissues, a classic and widely adopted technique uses a combination of Alizarin red for bone and Alcian or toluidine blue stains for cartilage. Published protocols for these stains have generally been optimized for small or prenatal specimens; thus, leaving out applications to relatively large specimens including somatically mature specimens and embryos of large-bodied taxa. Here, we present a newly refined, detailed protocol, accompanied with photographs, for clearing extraneous tissue and staining bone and cartilaginous structures optimized for larger avian specimens, including adult domestic chicken (*Gallus gallus domesticus*) and late-stage, embryonic emu (*Dromaius novaehollandiae*). The latter exhibit relatively mature soft tissue, including skeletal muscles, compared to those of late-stage embryos of small vertebrate taxa such as domestic chicken and mice. We found through our experiments that the combined Alcian blue and Alizarin red staining works best on freshly frozen specimens compared to those fixed in formalin and effective clearing and staining was possible without the use of tissue digestion reagents, such as trypsin and pancretin. The central aim of our work is to offer a step-by-step guide that provides researchers to consistently and effectively stain skeletal tissues in larger vertebrate specimens, allowing investigation of skeletal tissues at broader developmental and taxonomic scales.

## Introduction

Tissue staining is an indispensable tool for biological research, as it allows for powerful visualization of anatomical and molecular features of biological tissue. There are multiple types of staining protocols, such as those that utilize dyes (e.g., hematoxylin & eosin, Fischer et al., 2008; 4’,6-diamidino-2-phenylindole or DAPI, Kapuscinski, 1995), antibodies (Beutner, 1961), and radio-dense agents (Lugol’s iodine, Metscher, 2009; Gignac et al., 2016). These staining reagents each target a specific anatomical structure at a range of different scales. A classic and widely adopted staining method uses Alcian blue and Alizarin red dyes to stain cartilaginous and bone tissues, respectively, and then clearing the muscle tissues to easily visualize these internal structures (Spalteholz, 1914; Paysan, 2021). The combined usage of Alcian blue and Alizarin red staining has been widely adopted due to its utility for distinguishing between bone and cartilage, as well as between endochondral and dermal bones. Various protocols have been published for these stains (e.g., Rigueur & Lyons, 2014; Mead, 2020; Liao, et al., 2020). This procedure has been used to investigate a range of biological questions, including the effect of muscle paralysis on skeletal development (Rot-Nikcevic et al., 2006) and the repaired caudal skeleton from tail autonomy in geckos (Rakhmiyati & Jaâ, 2018).

To provide an overview of this staining technique: Alcian blue, a cationic dye, bonds to negatively charged cellulose in cartilage, whereas Alizarin red is an anionic dye that bonds to the positively charged fibers within the bone tissue (Rigueur & Lyons, 2014). A combination of alkaline solution (typically potassium hydroxide) and glycerol are then used to clear the surrounding tissues to visualize skeletal structures. This method of clearing during and after staining was pioneered by the anatomist Werner Spalteholz more than 100 years ago (Spalteholz, 1914), and modern organic solvent-based optical clearing techniques are derived from his original protocol. They all aim at creating a transparent sample through three basic steps: 1) removal of water (dehydration); 2) removal of lipids (delipidation); and 3) matching the refractive index to the average refractive index of the remaining constituents of the specimen, thereby creating a transparent sample. While dehydration is usually achieved by an alcohol gradient, delipidation and matching refractive index are both achieved by submerging the specimens with organic solvents such as potassium hydroxide (KOH) and glycerol. This last step requires careful consideration of the individual specimen being prepared for transparency to be successfully achieved (Paysan, 2021).

Although multiple protocols have been published for these stains (e.g., Rigueur & Lyons, 2014; Mead, 2020; Liao, et al., 2020), these dual bone and cartilage staining method has been used predominantly in small specimens, including adult individuals of small taxa and pre- or perinatal specimens (e.g., Sadeghi, 2014). In addition, most protocols have been established primarily in rodent models and have not been broadly tested on other taxa. While few exceptions exist with protocols on zebra fish, immature crocodylians and birds, seahorses and quail chicks (Witmer et al., 1995; Reed et al., 2019), they are not optimized for larger and more mature vertebrate specimens. While a study exists that features Alcian blue and Alizarin red stain on a late-stage emu embryo with relatively mature tissues (Nagai et al., 2011), it lacks a description of the protocol used by the authors. The clearing and staining technique of larger or more mature specimens present unique challenges due to differences in scale as well as properties of tissues. In fact, the motivation of this study originated from unsuccessful initial attempts at staining late-stage embryos of large, flightless ratite birds. While it is difficult to identify precisely which factors are contributing to the staining inconsistency in specimens with different stages of development, increased tissue density and composition discrepancies (e.g., proportions of aqueous matter, lipids, and proteins) are expected to reduce permeability of the staining solutions and thus influence the efficacy of the staining protocol. Taken together, there is a need for establishing a skeletal staining protocol that is optimized for larger and more mature avian specimens. to allow researchers to more effectively address biological questions requiring mature or large-bodied specimens.

To fill this gap, we present our newly refined protocol for clearing and staining relatively large avian specimens with well-developed tissues. We have optimized the protocol for two taxa and two anatomical regions: 1) the head of adult domestic chicken (*Gallus gallus domesticus*) and late-stage embryo of emu (*Dromaius novaehollandiae*); and 2) whole body of an adult chicken. The protocol below indicates when staining requirements differ (i.e., staining whole embryos at once can lead to over-staining of the head and under-stained postcrania). Furthermore, we also provide recommendations based on successful and unsuccessful trials. Our aim is to present a ‘recipe’ that investigators could follow to clear and stain larger and more mature specimens which could be challenging to achieve with existing protocols optimized for embryonic and smaller vertebrate specimens.

## Material and Methods

### Specimens

We sampled two heads from somatically mature (‘adult’) domestic chickens (*Gallus gallus domesticus*; Cornish cross Bovan breed) and late-stage embryos of the emu (*Dromaius novaehollandiae*). The chickens were sourced from Moody Farms (Poestenkill, New York, U.S.A.), and the emu specimens were collected from Boucher Family Farms (Firestone, Colorado, U.S.A.) (Table 1). For this study, the heads were prepared through *post-mortem* removal. Since these individuals died from natural causes, the study did not require IACUC approval. The specimens were initially frozen and were treated with different tissue fixation modes prior to the clearing and staining protocol: thawed from frozen state, fixed in 10% neutral buffered formalin, or fixed in graded ethanol from 25%, 50%, 75%, then 100%). The different types of tissue fixation allowed for investigation into which mode produces the best clearing and staining.

**Table 1.**
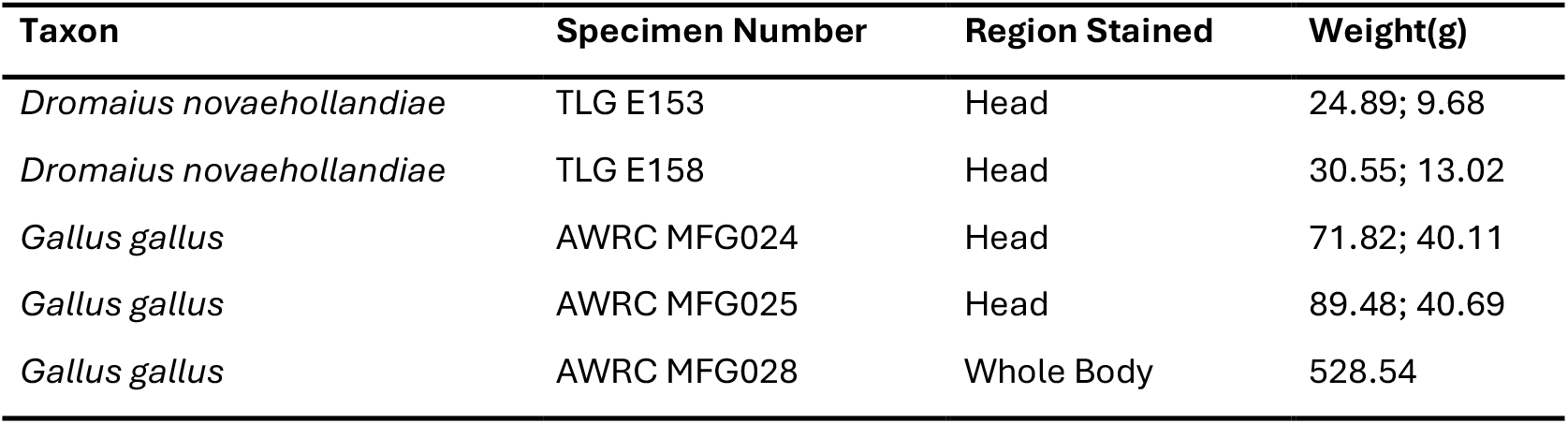
List of specimens sampled for this study. Specimen number refers to internal specimen catalog numbers used by the investigators (AWRC MFG: Akinobu Watanabe Research Collection Moody Farm Gallus specimens; TLG: Todd L. Green Research Collection). The weight values indicate the weight of the entire body for AWRC MFG028, and weight of the head with eyes and brain followed by weight of the head with eyes and brain removed for specimens TLG E153, TLG E158, AWRC MFG024, and AWRC MFG025.

| Taxon | Specimen Number | Region Stained | Weight(g) |
| --- | --- | --- | --- |
| *Dromaius novaehollandiae* | TLG E153 | Head | 24.89; 9.68 |
| *Dromaius novaehollandiae* | TLG E158 | Head | 30.55; 13.02 |
| *Gallus gallus* | AWRC MFG024 | Head | 71.82; 40.11 |
| *Gallus gallus* | AWRC MFG025 | Head | 89.48; 40.69 |
| *Gallus gallus* | AWRC MFG028 | Whole Body | 528.54 |

### Equipment

Besides basic dissection tools and glassware (e.g., 1000 mL beaker and graduate cylinder), we utilized the Benchmark Scientific Orbi-Shaker Low Speed Orbital Shakers for more even and efficient staining of samples and glass or high-density polyethylene (HDPE) containers that are large enough to fit the specimen(s) being stained (Table 2). Each container should be large enough to hold the specimen(s) without contacting the sides and with enough space to cover the specimen by 2.5–5 cm of solutions. We also used disposable transfer pipettes for removing the brain and gently cleaning out the eye and surrounding tissues in the orbit.

**Table 2.**
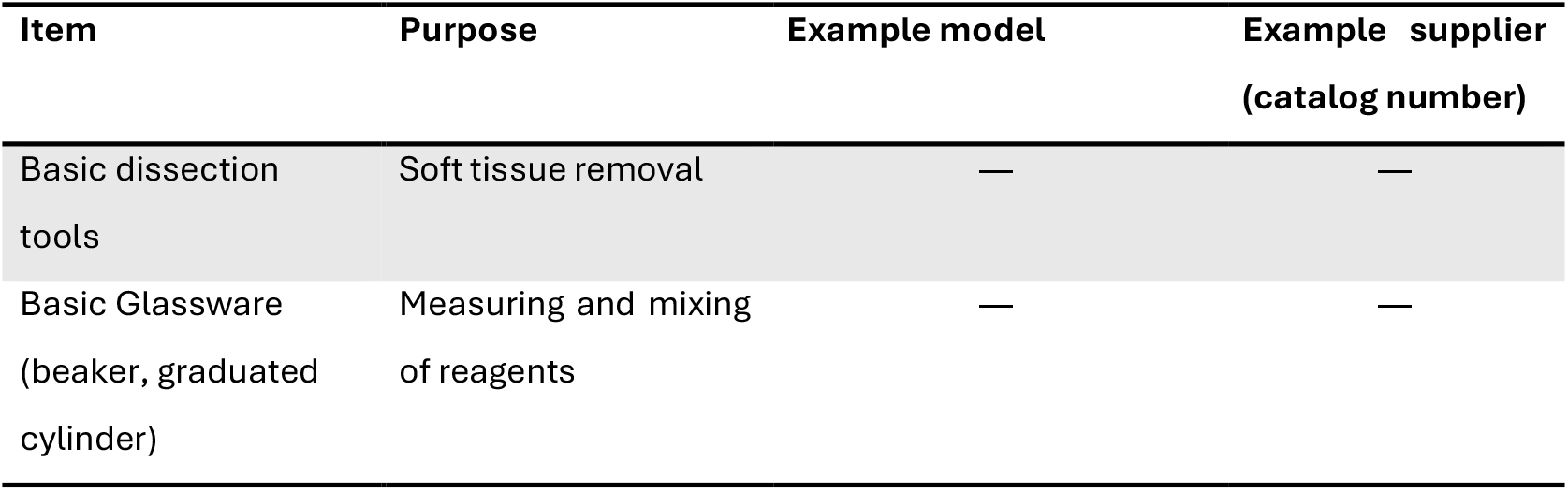
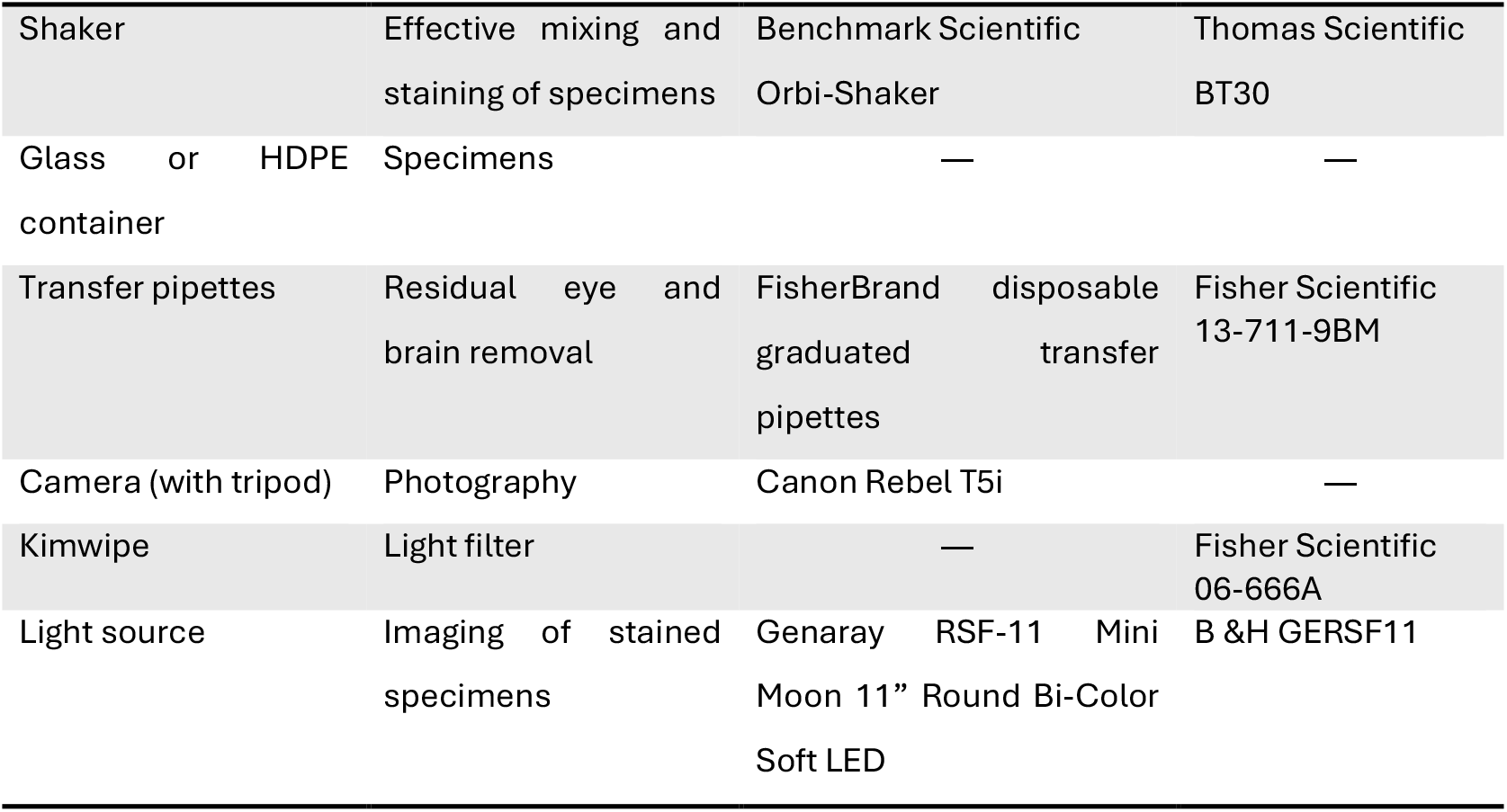
Equipment recommended for performing the clearing and staining protocol. We list specific models used in the study and their suppliers as suggestions.

### Imaging

To document each step of the protocol, we used an EOS Rebel T5i camera (Canon, Tokyo, Japan) with EF-S 18-135mm f/3.5-5.6 lens mounted on Alta series 60 tripod (Vanguard USA Inc., Michigan, U.S.A.). Specimens were photographed with the following settings: 1-second shutter speed, F18, and manual focus. ISO was set to ‘Auto’ except for photographing the specimen at the completion of the protocol, where it was set to 100. For imaging the completed specimen, we placed the RSF-11 Mini Moon 11-inch Round Bicolor Soft LED light (Genaray, New York, U.S.A.) to generate adequate amount of light to penetrate through the specimen. To reduce overexposure from bright lights at edges of the specimens, Kimwipes (Kimtech, Georgia, U.S.A.) were positioned around the periphery of the specimen to filter out some of the light.

### Reagents

We used 0.005% w/v Alizarin red solution for staining bone, which works best in basic and aqueous solution of potassium hydroxide (Table 3). We used 0.015% w/v Alcian blue solution for staining cartilage, which requires acidic and anhydrous environment to bind with mucopolysaccharides within the extracellular matrix of hyaline cartilage. Common laboratory chemicals were also used, including ethanol (200-proof to dilute appropriately) for fixation and dehydration of tissues; acetone, for lipid dissolution; glycerol, used for clearing and preservation of stained tissues, and potassium hydroxide solution, used to hydrolyze tissues (Table 3). We used phenol crystals to preserve the stored samples as well as inhibit mold growth.

**Table 3.** List of reagents used in the protocol and source. The ‘amount’ column indicates the amount of reagent typically needed for clearing and staining one whole-body chicken specimen.

| Reagent | Amount (whole-body stain) | Supplier (Catalog Number) |
| --- | --- | --- |
| Acetone | ~4L | Fisher Scientific (A18P-4) |
| Alcian blue 8GX | <1g | Fisher Scientific (AAJ6012209) |
| Alizarin red | <1g | Fisher Scientific (AC400480250) |
| Ethanol (200-proof) | ~4L (for whole-body) | Greenfield Global (100129) |
| Glacial Acetic Acid | ~4L (for whole-body) | Fisher Scientific (LC1010036) |
| Glycerol | ~4L (for whole-body) | BioBasic (GB0232) |
| Phenol (Crystals) | Few crystals | Fisher Scientific (C956M52) |
| 30% (w/v) Potassium Hydroxide | ~1L for whole-body | Fisher Scientific (6153-32) |

## RESULTS

### Protocol

This protocol assumes beginning with a whole avian specimen. After each step, solutions and specimens should be left on a shaker at room temperature. For staining the head and whole-body specimens, one should expect the protocol to take a total of 21 and 47 days, respectively. Please refer to Figure 1 and Table 4 for an overview of the steps and duration of each.

**Table 4.**
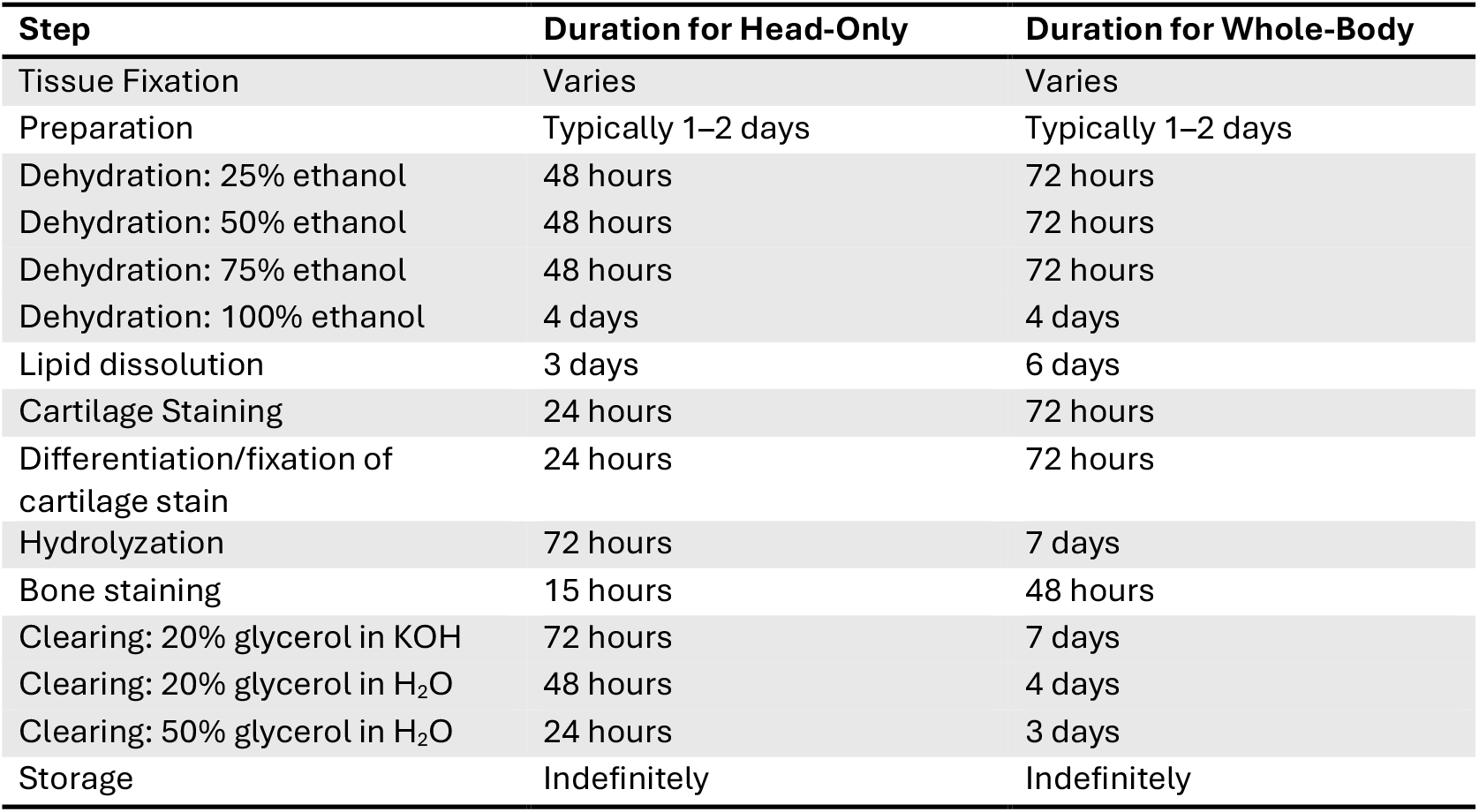
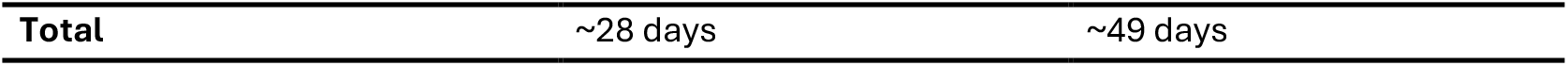
Duration of each step of dual staining of bone and cartilage for head-only and whole-body specimens.

**Fig. 1.**
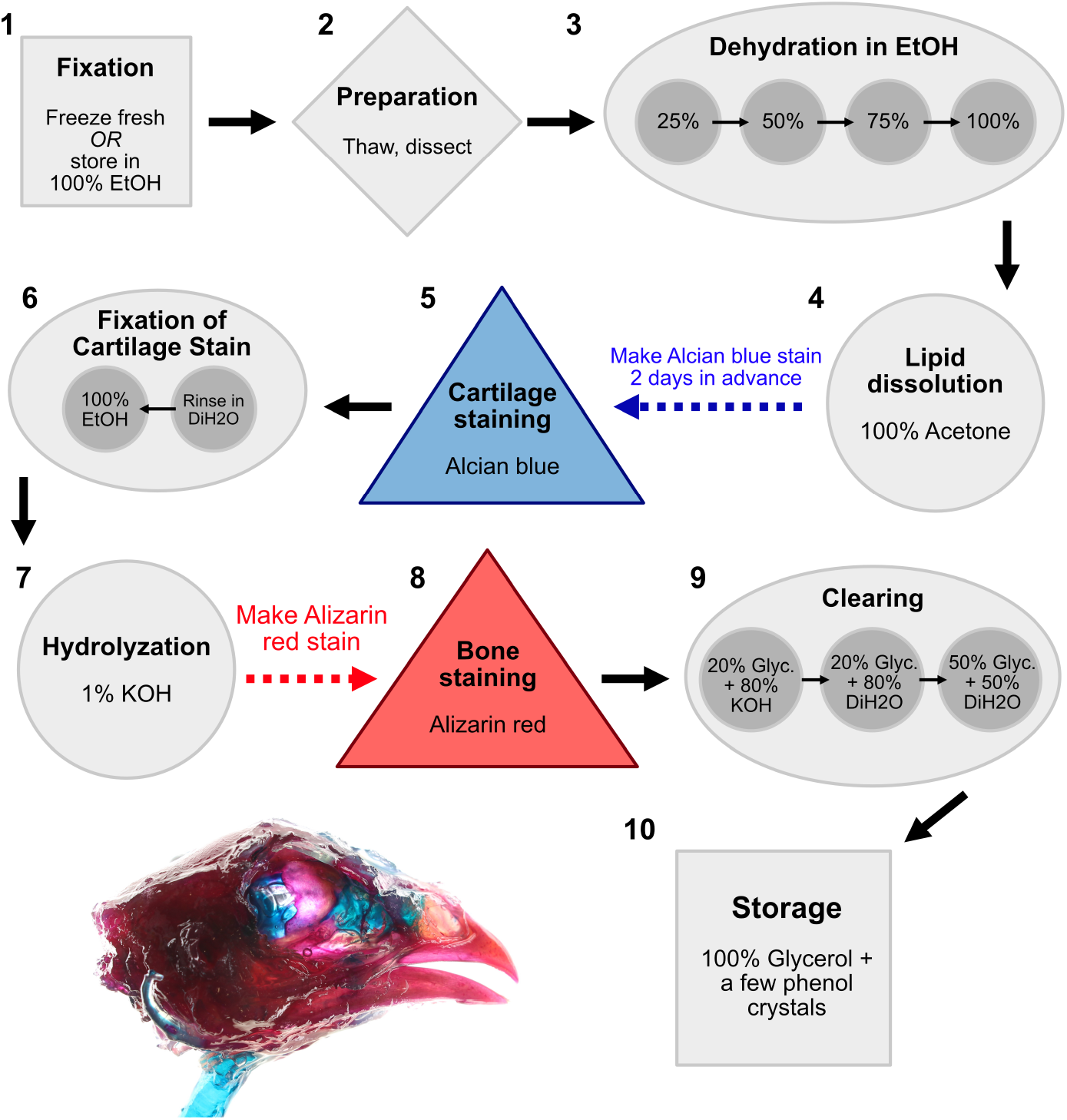
Schematic figure of protocol pipeline, with numbers corresponding with protocol steps. Shapes indicate the task type for each step; Square = storage; Circle = rinsing; Triangle = staining; Diamond = miscellaneous. Durations for each step vary depending on the specimen size and whether the head or whole body is being stained (see Results: Protocol). Specimen photo is a head of *Gallus gallus* (AWRC MFG 025) at the completion of the protocol.

1. **Tissue Fixation:** This procedure works best if the specimens are frozen fresh and thawed immediately before beginning protocol. Alternatively, specimens can be stored in 100% (200-proof) ethanol if needed. We advise against fixing tissues in 10% neutral-buffered formalin or other formaldehyde-based fixatives because this can lead to Alcian blue stain remaining within the tissues of the specimens even after clearing steps are conducted (discussed below).
2. **Preparation**: If specimens were frozen, they should be thawed on benchtop at room temperature for ∼3 hours during the day or overnight in a refrigerator. Thawed avian specimens should be soaked in a glass or HDPE container with warm tap water (up to 55º C) for 3-5 minutes to loosen skin. Thawing should be conducted sufficiently to allow for soft tissue to be dissected and removed. Skin and feathers should be removed carefully. For whole-body staining, the scaly integument (e.g., feet) should be removed to allow for stains to penetrate underlying tissues effectively. Eyes are removed from the orbits using dissection tools and disposable transfer pipettes to remove remaining tissue; however, the intact eyes with sclerotic ossicles can be included with the specimen to be cleared and stained if interested. In addition, brain tissue should be removed from skull cavity through the foramen magnum using distilled water and disposable transfer pipettes. If only the head is being stained, then the head should be removed at mid-cervical series using dissection tools. For whole-body staining, one should ensure that as much of the soft tissue is removed, including muscles, viscera, and adipose tissue. We recommend keeping the trachea intact, even if the structure is not of direct interest to one’s study, as it serves as good reference for cartilage staining. The specimen should be placed into a glass or HDPE container that comfortably fits the specimen— approximately twice the volume of the specimens for sufficient submersion. All solutions through each step should cover the top of the specimen by 2.5 - 5.0 cm.
3. **Dehydration**: The specimens should pass through graded ethanol baths of increasing concentrations until 100% ethanol is achieved. We submerged the specimen with 25%, 50% and 75% concentrations, and the specimen should stay at each concentration of ethanol for 48 hours for head-only, and 72 hours for whole-body specimens. For larger, whole-body adult specimens, one should refresh the solution on the second of the three-day stages at each ethanol concentration. Once 100% ethanol concentration is reached, the specimen should be submerged in the solution for 96 hours (4 days) or more. The tissue should appear pale and feel tough to the touch after dehydration (Figs. 2a, 3a, 4a).
4. **Lipid Dissolution**: The specimen is removed from ethanol and put in 100% acetone for 72 hours (3 days) and 6 days for whole-body specimen. This process may cause some minor shrinkage of the tissues (Figs. 2b, 3b, 4b). During this step, one should prepare the Alcian blue solution two days before proceeding to cartilage staining (see next step for instructions).
5. **Cartilage Staining:** The staining solution should be prepared two days ahead of time. We found that cartilage staining is more consistent and stable against over-leaching if the staining solution is made ahead of time. We prepared 0.015% w/v Alcian blue stain solution for both head and whole-body staining, which has the following composition: 80% of 200-proof ethanol, 20% of glacial acetic acid, and 0.015% of Alcian blue powder. This should be mixed well and left on counter until used on a specimen. For each 100 mL of Alcian blue solution, we used 0.015g of powder, 80 mL of 100% ethanol (200 proof), and 20 mL of glacial acetic acid. For a 4.5 L stock, one can combine 3600 mL of 100% ethanol, 900 mL of glacial acetic acid, and 0.675 g of Alcian blue 8GX powder. After acetone submersion, the Alcian blue solution is agitated vigorously, then poured into the specimen container after draining acetone (this reagent can be reused a few times). Specimens can be left in the Alcian blue solution for 24 hours for the head-only specimen and 72 hours for the whole-body specimen. Specimens will be strikingly blue when removed, and all parts of the specimen will be almost completely stained (Figs. 2c, 3c, 4c). The excess of the stain will be removed from non-cartilage tissues in successive steps.
6. **Differentiation/Fixation of Cartilage Stain:** The specimen is rinsed under running (distilled) water and transferred into 100% ethanol, then left for 24 hours for head-only and 72 hours for whole-body specimens. This will begin to remove the Alcian blue stain from the non-cartilaginous tissues.
7. **Hydrolyzation:** The specimen is then placed into 1% potassium hydroxide solution (prepared with distilled water for 72 hours), and changed to fresh solution as the solution becomes blue as Alcian blue stain is removed from the specimen. For a whole-body stain, this process could take up to 7 days or longer depending on size until the potassium hydroxide solution remains mostly clear. The solution should be replaced until blue color is no longer leaching into the solution, although some blue stain may still be on non-cartilaginous portions of the specimen (Figs. 2d, 3d, 4d). As soon as the solution remains clear, one can proceed to the next step. However, overexposure to potassium hydroxide solution will lead to disintegration of the specimen. As such, it is important to consider the delicate trade-off between de-staining of Alcian blue stain and keeping the specimen intact. During the hydrolyzation and two days prior to proceeding to the next step, the Alizarin red solution should be prepared (see next step for instructions).
8. **Bone Staining:** Next, the specimen is transferred to a 0.005% w/v Alizarin red solution (prepared in 1% potassium hydroxide solution two days prior to proceeding to the Alizarin red staining) for 15 hours for the head-only and 48 hours for the whole-body staining (Figs. 2e, 3e, 4e). For each 100 mL of Alizarin red solution, we used 0.005 g of Alizarin red powder and100 mL of 1% potassium hydroxide solution in distilled water. More practically, one could create a 4 L solution by combining 4000 mL of 1% potassium hydroxide with 0.2 g of Alizarin red powder. The stain may appear purple than “red” as implied by the reagent name, but this is normal.
9. **Clearing:** The specimen is then rinsed in distilled water and transferred into clearing solution at the end of the alizarin red staining. The following steps will finish the clearing process.
  - The specimen should be left in a glycerol and potassium hydroxide (2%) solution. Each 100 mL solution comprises 20 mL of glycerol and 80 mL of 2% KOH solution. The solution should be replaced each day for a head-only specimen and every 3–4 days for a whole-body specimen until the color from the Alizarin red and Alcian blue stains stops leaching. This could take at least 72 hours and 7 days for whole-body specimens.
  - Then, place the specimen in a 20% glycerol solution. Each 100 mL solution comprises 20 mL of glycerol with 80 mL of distilled water, and the specimen should be left in the solution for 48 hours for head-only and 4 days for whole-body specimens.
  - Finally, the specimen is placed in a 50% glycerol solution. Each100 mL solution comprises 50 mL glycerol with 50mL of distilled water, and the specimen should be left in the solution for 24 hours for head-only and 3 days for whole-body clearing. By the end of the clearing steps, the soft tissue of the specimen should be opaque with consistent blue and red staining on cartilage and bone (Fig. 2f; 3f; 4f).
10. **10. Storage:** The specimen is then transferred to a glass container in 100% glycerol. Approximately three phenol crystals are to be added for long term storage.

**Fig. 2.**
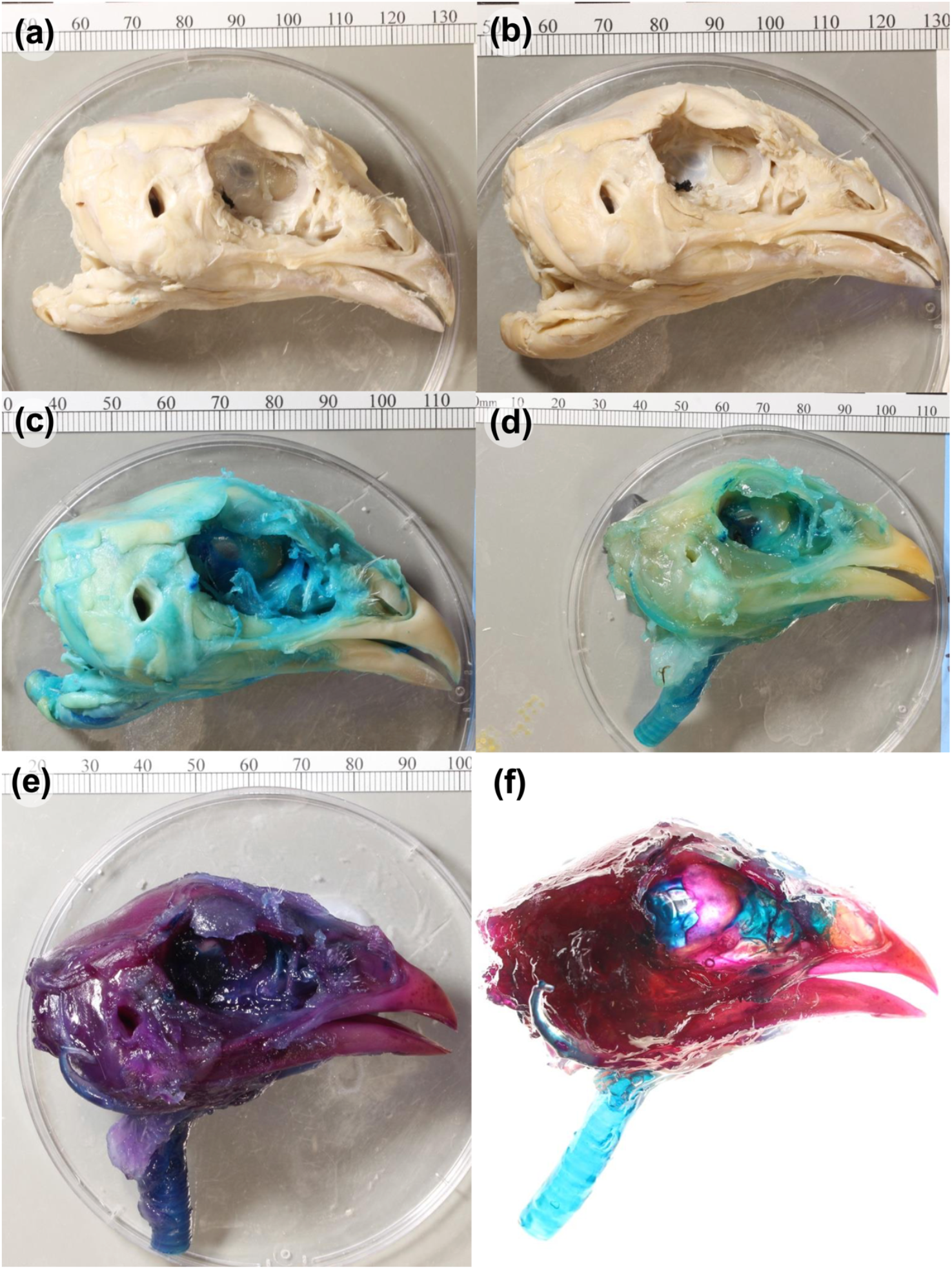
Photographs of a chicken head (AWRC MFG025) at select stages of the clearing and staining protocol in right lateral view after (a) soft-tissue removal and dehydration; (b) lipid dissolution step; (c) Alcian blue stain; (d) differentiation and fixation of cartilage stain and hydrolyzation; (e) Alizarin red stain; and (f) completion of clearing and staining protocol.

**Fig. 3.**
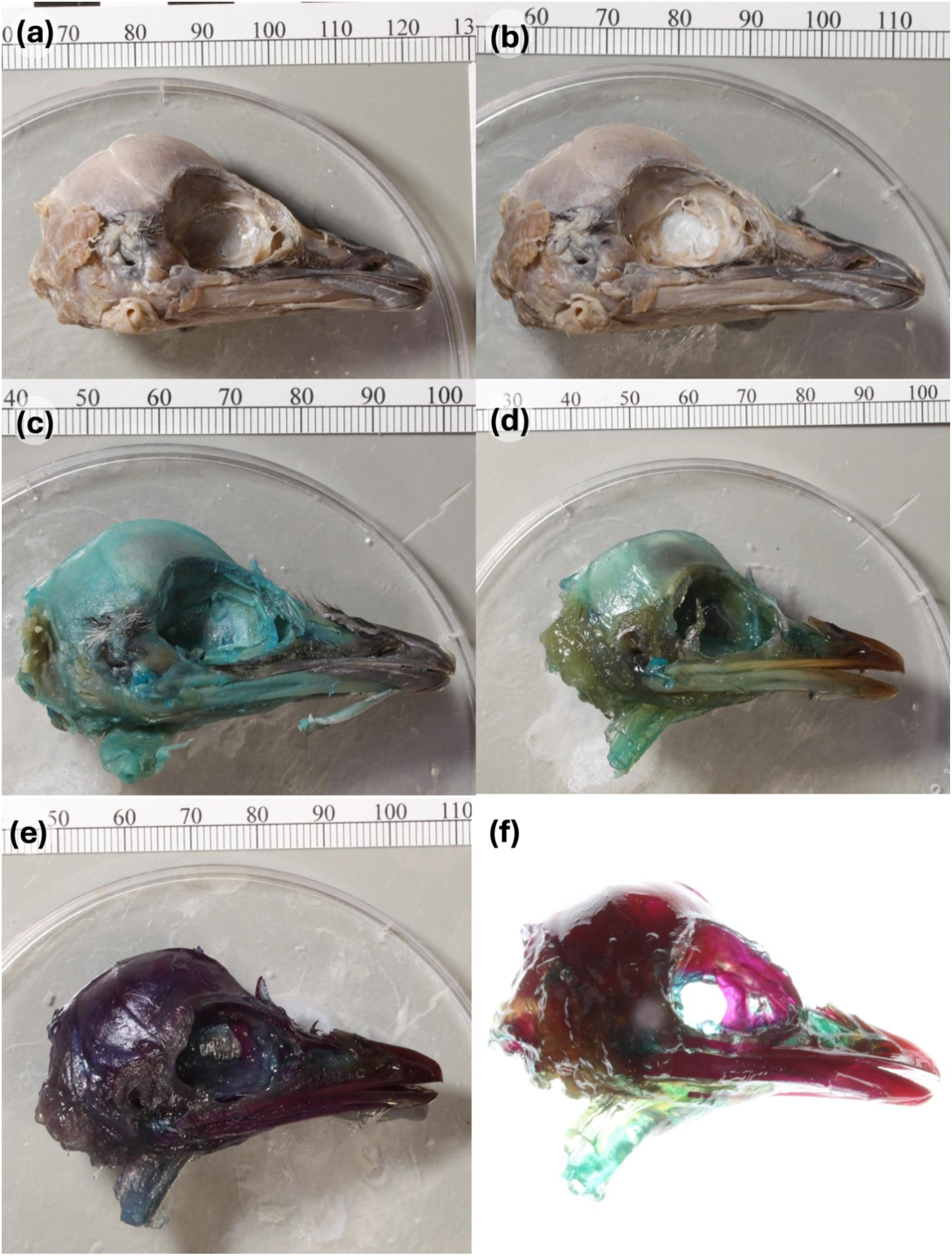
Photographs of an emu head (TLG E158) at select stages of the clearing and staining protocol in right lateral view after (a) soft-tissue removal and dehydration; (b) lipid dissolution step; (c) Alcian blue stain; (d) differentiation and fixation of cartilage stain and hydrolyzation; (e) Alizarin red stain; and (f) completion of clearing and staining protocol.

**Fig. 4.**
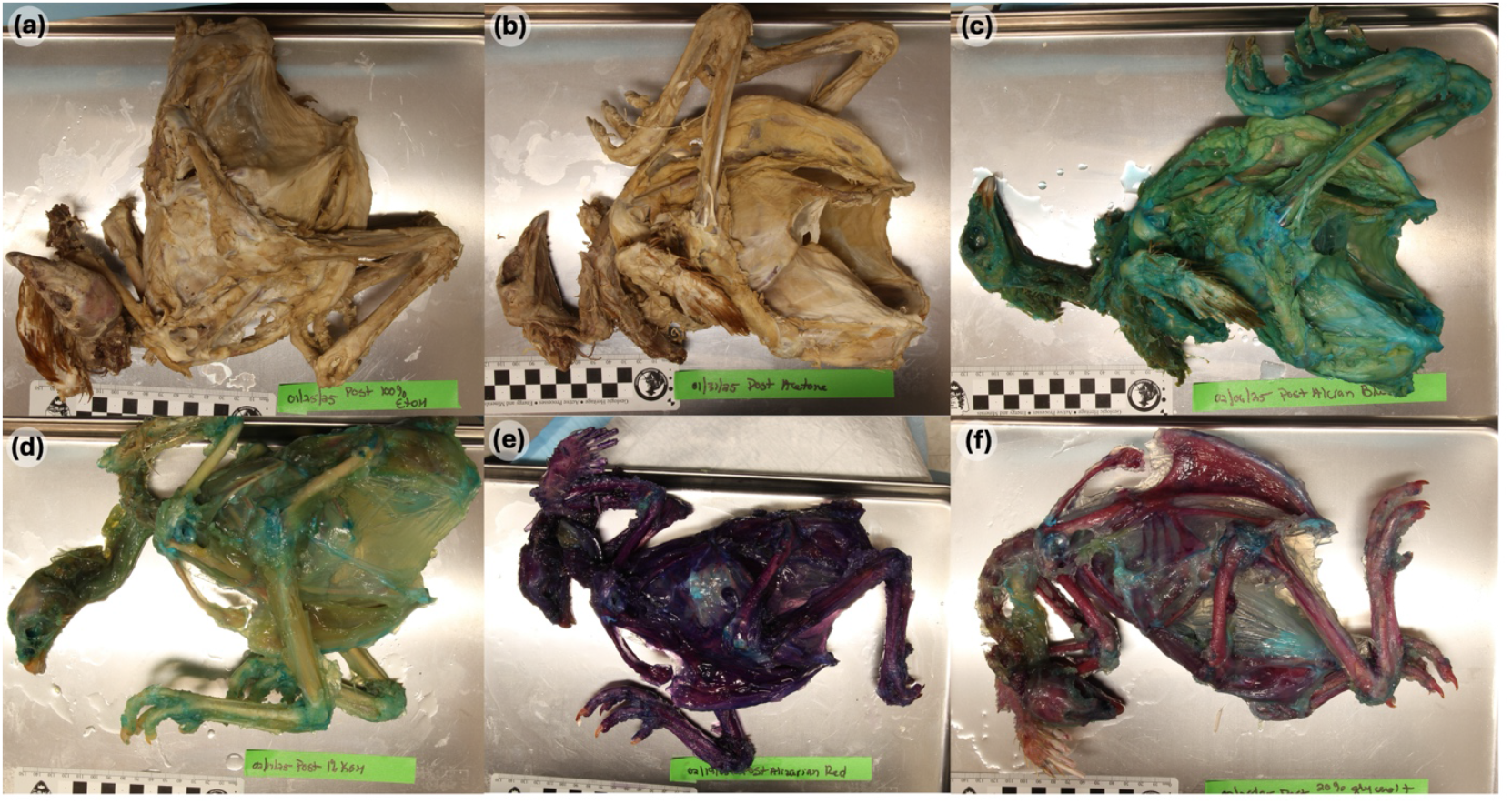
Photographs of whole-body domestic chicken (AWRC MFG028) after (a) soft-tissue removal and dehydration; (b) lipid dissolution step; (c) Alcian blue stain; (d) differentiation and fixation of cartilage stain and hydrolyzation; (e) Alizarin red stain; and (f) completion of clearing and staining protocol.

## Discussion

The protocol presented above represents a step-by-step instructional workflow based on successful and unsuccessful trials of clearing and staining of larger avian specimens than reported in other protocols on bone and cartilage staining in vertebrates. In this section, we present additional insights on the best practices for clearing and staining these classes of specimens, particularly in tissue removal, fixation, and clearing procedures.

### Tissue Removal

We found that careful, but liberal removal of soft tissue is necessary for effective clearing and staining of large, more mature specimens. In our initial attempts, we performed a relatively conservative removal of soft tissue for the whole-body staining, where much of skeletal muscle was kept on the specimen. This led to difficulty in the stain, especially Alcian blue stain, to penetrate to their target tissue type. Moreover, the blue stain generally remained on skeletal tissues even after the clearing procedure. Therefore, we recommend that as much soft tissue is removed from the specimen as possible while keeping the integrity of the specimen.

### Fixation

We found that the initial mode of tissue preservation is critical in determining the quality of clearing and staining. Our initial attempt to use formalin-fixed specimens led to difficulty in clearing off the Alcian blue stain (Fig. 5). While we did not investigate the mechanistic explanation for this observation, formalin and other formaldehyde-based solutions crosslinks proteins and/or nucleic acids in the tissue by reacting with their amine groups, and this preservation mode may cause the non-specific binding of Alcian blue stain. This is an intriguing observation because formaldehyde-based fixatives have been successfully used in published bone and cartilage stain protocol of embryonic (e.g., Carneiro et al., 2021) and small or regional specimens (e.g., Lima et al., 2017; Tsandev et al., 2017). Therefore, this issue with use of formaldehyde-based fixative may only apply to relatively mature specimens with more developed soft tissues. Over-fixation of tissues with paraformaldehyde is known to result in lower quality fluorescent labelling from fixative molecules blocking access to the epitopes than ethanol fixation (Chung et al., 2018; Richardson et. al., 2021). Similarly, formaldehyde-based fixation could alter the affinity of both cartilaginous and non-cartilaginous tissues to Alcian blue, preventing the stain from being specific or be removed through the latter clearing steps. In contrast, initial long-term fixation with ethanol, which preserves tissues through dehydration of proteins, showed diminished but adequate clearing and staining (MJT pers. obs.). Taken together, we recommend using frozen specimens and minimal use of tissue fixation methods prior to the first dehydration step of the clearing and staining protocol.

**Fig. 5.**
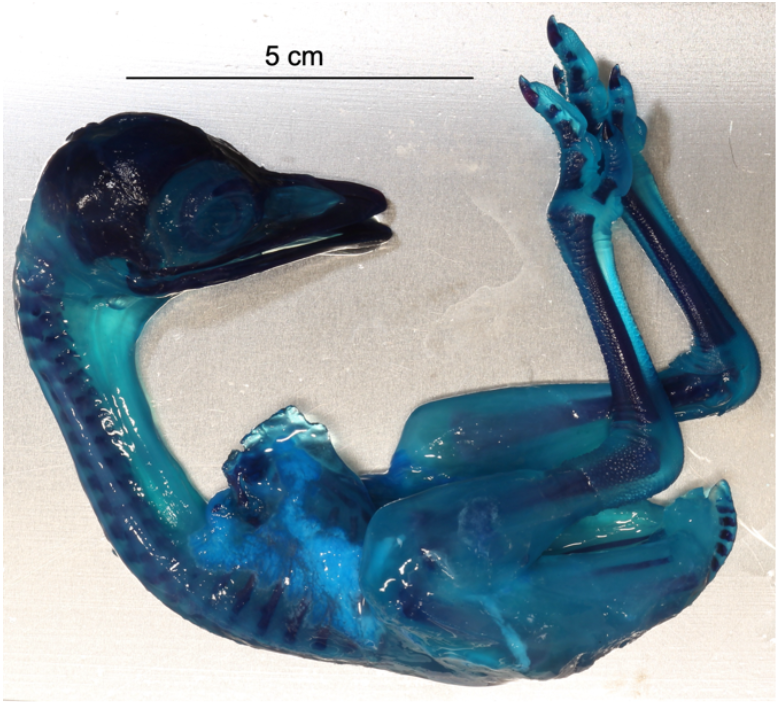
Initial attempts at clearing and staining embryonic emu specimen (TLG E139, not formerly included as sample in this study) fixed in 10% neutral-buffered formalin. Note that the Alcian blue stain remained on non-cartilaginous tissues despite completion of the initial version of the protocol.

### Clearing

One of the most difficult aspects of the clearing and staining process was removing the excess Alcian blue stain from the non-cartilaginous tissues in the birds. This issue may have been caused by formalin-based fixatives as described above, the size and density of the muscle tissues related to their maturity, or likely both. We tried various methods based on other published protocols including using a trypsin or pancretin solution to help breakdown remaining skeletal muscle tissue, using a saturated sodium borate solution to help remove and prevent the Alcian blue from binding to non-cartilaginous tissues, and using hydrogen peroxide as a bleaching agent to remove stain. Tissue digestion, nor bleaching agents such as hydrogen peroxide were not necessary when staining the crania nor whole-body specimens. We also tested varying duration for each step of the staining and clearing process, ranging from 30 minutes of graded ethanol stages to 40 days of trypsin submersion. We found that using graded ethanol steps increased the binding capacity of the stains, compared to immediately putting the samples directly into 100% ethanol. Through trial and error, we found that making the stains a few days ahead of time before using on samples for all powder to dissolve completely in solution. Therefore, using frozen specimens, graduated ethanol, making stains ahead of time, and carefully monitoring the time spent in potassium hydroxide solution, were enough to successfully clear and stain the adult chicken and embryonic emu heads.

## Conclusions

Here, we present a protocol for bone and cartilage staining that has been optimized for heads and whole-body specimens of large, late-stage emu embryos and somatically mature chickens. Our goal is to provide a clear, detailed, and ready-to-use guide to apply dual bone-cartilage staining to heads and whole bodies of vertebrate taxa. Being able to differentiate and identify cartilaginous and bony tissues *in situ* in large-bodied embryonic and mature avian specimens opens doors for studies of internal structures to include greater taxonomic and developmental diversity than was previously possible with previously published protocols.

## Author Contributions

MJT: Methodology, investigation, writing—original draft. TLG: methodology, investigation, writing— review and editing. GG, TL-M: investigation, writing—review and editing. AW: conceptualization, methodology, investigation, writing—review and editing.

## Acknowledgments

We would like to sincerely thank Nathan Kley (Stony Brook University), Lawrence Witmer (Ohio University), and Christian Louis Bonatto Paese (Allen Institute) for sharing their protocols and discussion that formed the initial template and guided our development of the protocol; Allison Arnold (New York Institute of Technology) for processing orders of specimens and reagents; and Rachel Moody (Moody Farms) and Lisa Nowak (University of Connecticut Poultry Farm), and Boucher Family Farms for providing the avian specimens sampled in this study.

## Notes

**Conflict of Interest** Statement The authors declare no conflict of interest.

### Competing Interest Statement

The authors have declared no competing interest.

